# Mating in wild yeast: variable interest in sex after spore germination

**DOI:** 10.1101/360289

**Authors:** Allison W. McClure, Katherine C. Jacobs, Trevin R. Zyla, Daniel J. Lew

## Abstract

Studies of lab strains of *Saccharomyces cerevisiae* have uncovered signaling pathways involved in mating, including information processing strategies to optimize decisions to mate or to bud. However, lab strains are heterothallic (unable to self-mate) while wild yeast are homothallic. And while mating of lab strains is studied using cycling haploid cells, mating of wild yeast is thought to involve germinating spores. Thus, it was unclear whether lab strategies would be appropriate in the wild. Here, we have investigated the behaviors of several yeast strains derived from wild isolates. Following germination, these strains displayed large differences in their propensity to mate or to enter the cell cycle. The variable interest in sex following germination was correlated with differences in pheromone production, which were due to both cis- and trans-acting factors. Our findings suggest that yeast spores germinating in the wild may often enter the cell cycle and form microcolonies prior to engaging in mating.

## Introduction

The molecular and genetic tractability of the budding yeast, *Saccharomyces cerevisiae*, has made it a premier model organism for the study of many aspects of biology (Botstein and Fink, 2011; Botstein et al., 1997). Among these, the yeast mating pathway has yielded paradigm-setting discoveries concerning signal transduction, pheromone biogenesis, regulation of gene expression, and the cell biology of chemotropism and cell-cell fusion (Arkowitz, 2009; Atay and Skotheim, 2017; Merlini et al., 2013; Michaelis and Barrowman, 2012). While molecular mechanisms have been well-studied in the laboratory, surprisingly little is known about *S. cerevisiae* mating outside the lab. Recent work has emphasized the importance of studying the life history of *S. cerevisiae* as well as other model organisms in order to properly interpret laboratory findings (Boynton and Greig, 2014; Knop, 2006; Liti, 2015). Here we consider ways that mating may differ in the lab and in the wild.

Lab strains of yeast can be grown as diploids, or they can be maintained as haploids of either mating type (MAT**a** or MATα). In the lab, mating assays are conducted under “orgy” conditions (Hartwell, 1973) in which large numbers of **a** and a cells first encounter each other when they are abruptly mixed together by the investigator. Haploids secrete small peptide pheromones, a-factor and α-factor, which trigger a number of changes to prepare cells for mating (Alvaro and Thorner, 2016). Successful mating involves arrest in G1 phase of the cell cycle, polarization of growth towards the mating partner, and expression of numerous genes that enhance cell-cell adhesion and eventual fusion of the cells and nuclei to form a diploid zygote.

Successful mating is not guaranteed: for example, the non-motile haploid cells may be too far apart, or another partner may mate with the intended target. Recent studies have revealed a sophisticated decision-making system that appears optimized for mating success. Cells in liquid media can optimize their decision to mate or to proliferate by detecting the ratio of opposite-sex partners to same-sex competitors (Banderas et al., 2016). Cells on solid media detecting sub-optimal pheromone levels undergo a “chemotropic growth” program in which they arrest transiently and grow towards the mating partner, but then re-enter the cell cycle and form a bud in the direction of growth (Erdman and Snyder, 2001; Hao et al., 2008). This response places the resulting daughter cell closer to the pheromone source, presumably increasing the chances that it will mate. In the following cell cycle, the cells retain and exploit a memory of the fact that they had previously responded to pheromone: the original “mother” cell no longer arrests in response to the same low levels of pheromone (Caudron and Barral, 2013), while the daughter cell is more likely to arrest in response to low pheromone levels (Doncic et al., 2015). Thus, cells with better chances of mating are primed to mate, while cells unlikely to mate invest their energies more productively in vegetative growth.

These decision-making processes were uncovered using lab strains, and the behaviors were interpreted to optimize outcomes under specific lab mating conditions. However, lab strains differ from wild strains in an important respect: the ability to switch mating type. After producing its first daughter, a haploid mother cell switches mating type in the next cell cycle, while its daughter does not (Haber, 2012). This means that the next cell cycle will yield cells of both mating types in close proximity, providing an opportunity to mate in a process termed “haplo-selfing” (Fig. 1A). Due to the presence of mating-type switching, wild yeast strains do not proliferate stably as haploids. Almost all wild *S. cerevisiae* isolates (environmental and clinical) are diploid.

**Figure 1.**
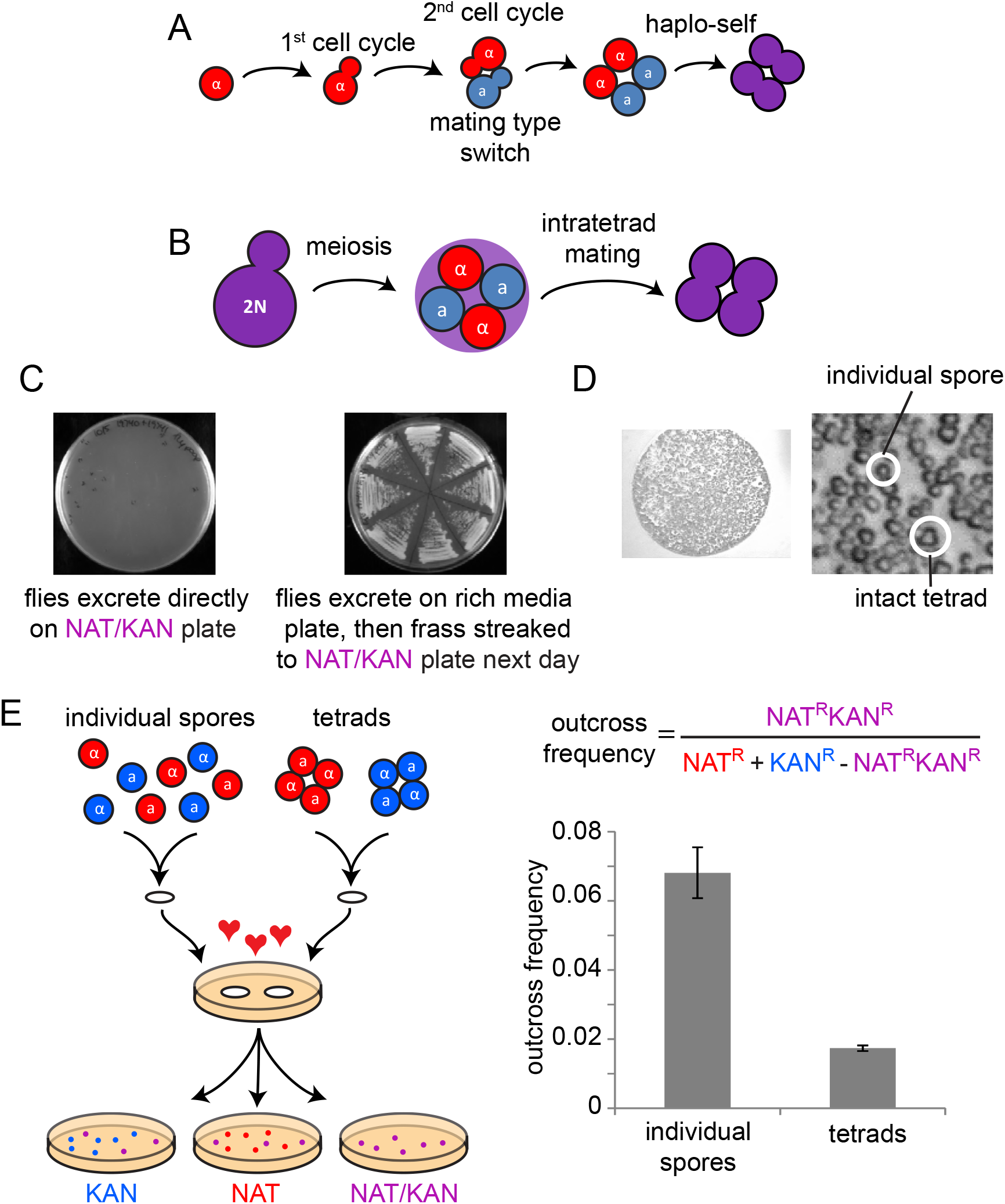
Yeast mating scenarios. (A) Cartoon depiction of mating type switching allowing for haplo-selfing. (B) Cartoon depiction of intratetrad mating. (C) Left: flies were fed tetrads that carried either NAT or KAN resistance markers (DLY19740 and DLY19741), and then allowed to excrete on a NAT/KAN double drug plate for 1 h. Excretion spots were marked on the cover of the plate, but showed no growth after 2 days. Right: after feeding, flies were allowed to excrete on a plate without drugs for 1 h. The next day, excretion spots showed yeast growth that was then streaked to a NAT/KAN double drug plate. (D) Excretion spots show a mix of intact tetrads and isolated spores. (E) Either individual spores or intact tetrads with drug resistance markers (indicated by blue or red) were placed on filters where cells germinated and mated overnight. Then, cells were recovered from the filters and plated to identify the rate of outcrossing. Separated spores showed a higher outcross frequency than intact tetrads (mean +/-SEM of three plates, t-test, p<0.006).

The circumstances under which mating occurs in the wild are not well understood. Unfavorable nutritional conditions initiate a program of meiosis and sporulation, whereby a diploid cell generates a tetrad with four haploid spores in an ascus (Neiman, 2011). Spores are metabolically dormant and have a tough outer cell wall; they can survive harsh conditions and prolonged starvation. Upon exposure to fermentable sugars, spores germinate, awaken metabolic pathways, swell, and break the outer cell wall (Herman and Rine, 1997; Joseph-Strauss et al., 2007). If a partner of the opposite mating type is available (e.g. a sibling spore in a tetrad), germinating spores can mate to regenerate a diploid (Taxis et al., 2005) (Fig. 1B). This “intratetrad mating” would differ from that examined in lab-based mating assays because the germinating spore has a different physiology from that of a haploid cell grown for many generations in rich media, and because there would be only one or two available partners to choose from within the tetrad.

If no suitable partner were present when a spore germinates, the haploid cell would enter the cell cycle, form a bud, and then switch mating type in the next cycle. Even if no mating partner is close by, mating-type switching guarantees that a partner will soon become available immediately adjacent to the cell. In that context, it is unclear whether the decision-making strategies uncovered for lab strains would make sense for wild strains of yeast.

To better understand the mating behaviors of wild yeast, we examined the events following germination of spores (either in tetrads or on their own) from various wild and lab strains. Depending on the strain, the frequency of intratetrad mating varied widely. For several strains, germinating spores and their progeny chose to bud instead of mate to a nearby potential partner, generating microcolonies with haploid cells of both sexes (due to mating-type switching). Mating subsequently took place in the microcolony context. Pheromone production upon germination was variable between strains, probably contributing to the variable interest in mating. Our findings suggest that mating in the wild may often occur in the context of mixed mating-type microcolonies, with implications for decision making and outbreeding.

## Results

### Potential mating scenarios for wild yeast

In the wild, insects are thought to provide dispersal vectors for yeast (Gilbert, 1980; Reuter et al., 2007). Insects carry yeast cells on their legs, and insects eat yeast cells. In a remarkable recent study, diploid yeast cells were fed to wasps that were then induced to hibernate. Over the ensuing weeks, the yeast underwent meiosis, sporulation, germination, and mating, all in the digestive tract of the hibernating wasp (Stefanini et al., 2016), providing one potential wild yeast mating scenario. Although diploid yeast cells would likely be digested following ingestion by active (non-hibernating) insects, yeast spores were shown to survive passage through the digestive tract of fruit flies (Coluccio et al., 2008; Reuter et al., 2007). Flies were also shown to increase the frequency of yeast outbreeding (Reuter et al., 2007), suggesting that spores that survive digestion might subsequently germinate and mate in fly frass. We fed *Drosophila melanogaster* flies tetrads from two strains of yeast, each carrying a different drug resistance marker. When the flies excreted directly onto solid media containing both drugs, no cells grew (Fig. 1C), indicating that no detectable mating took place within the gut of the fly. However, if the flies excreted on solid media lacking drugs, and the frass was streaked onto media containing both drugs the next day, many outbred colonies grew. This supports the idea that wild yeast might mate following spore germination in fly frass.

Although the flies were fed tetrads, digestive enzymes within the fly gut have been shown to separate sibling spores (Reuter et al., 2007). Imaging the fly frass, we found a mix of intact tetrads and isolated spores (Fig. 1D). It was suggested based on indirect evidence that isolated spores would be more likely than spores in tetrads to engage in outbreeding (Reuter et al., 2007). Consistent with that hypothesis, we found using a quantitative mating assay that non-sibling mating was more common when starting with isolated spores than with tetrads (Fig. 1E). These findings support the idea that in the wild, yeast spores or tetrads might be deposited in a new and nutrient-rich environment following excretion by flies, and that mating occurs following spore germination in fly frass. However, it was not clear whether the germinating spores constitute the cells that actually mated, or whether mating followed one or more haploid vegetative cycles.

### Mating behavior of germinating spores in tetrads

To study the mating behaviors of wild yeast strains, we selected a set of nine genetically diverse strains from the 100-genomes collection (Strope et al., 2015) (Fig. 2A), derived from isolates collected from different environments (Supplementary Table 1). These homothallic strains are homozygous diploids derived by mating-type switching and haplo-selfing. Diploids were incubated in potassium acetate media to induce sporulation, and tetrads were placed on microscope slabs with rich media and imaged by time-lapse microscopy to observe behaviors following germination. Upon germination in a tetrad, spores either mated with a sibling (Fig. 2B) or entered the cell cycle and formed a bud (Fig. 2C). Under these conditions, many germinating spores entered the cell cycle even if a suitable mating partner was present. Quantification of the percentage of spores that mated or budded following germination revealed that even in the wild strain with the highest propensity to mate, around 20% spores entered the cell cycle rather than mate (Fig. 2D). The mating behavior was variable between wild strains, but intratetrad mating was generally much lower than in a previously described lab strain (Taxis et al., 2005). This does not appear to represent a general difference between wild and lab strains, because two S288C-derived lab strains (labeled 15D and YEF) showed lower mating propensities than most of the wild strains (Fig. 2D).

**Figure 2.**
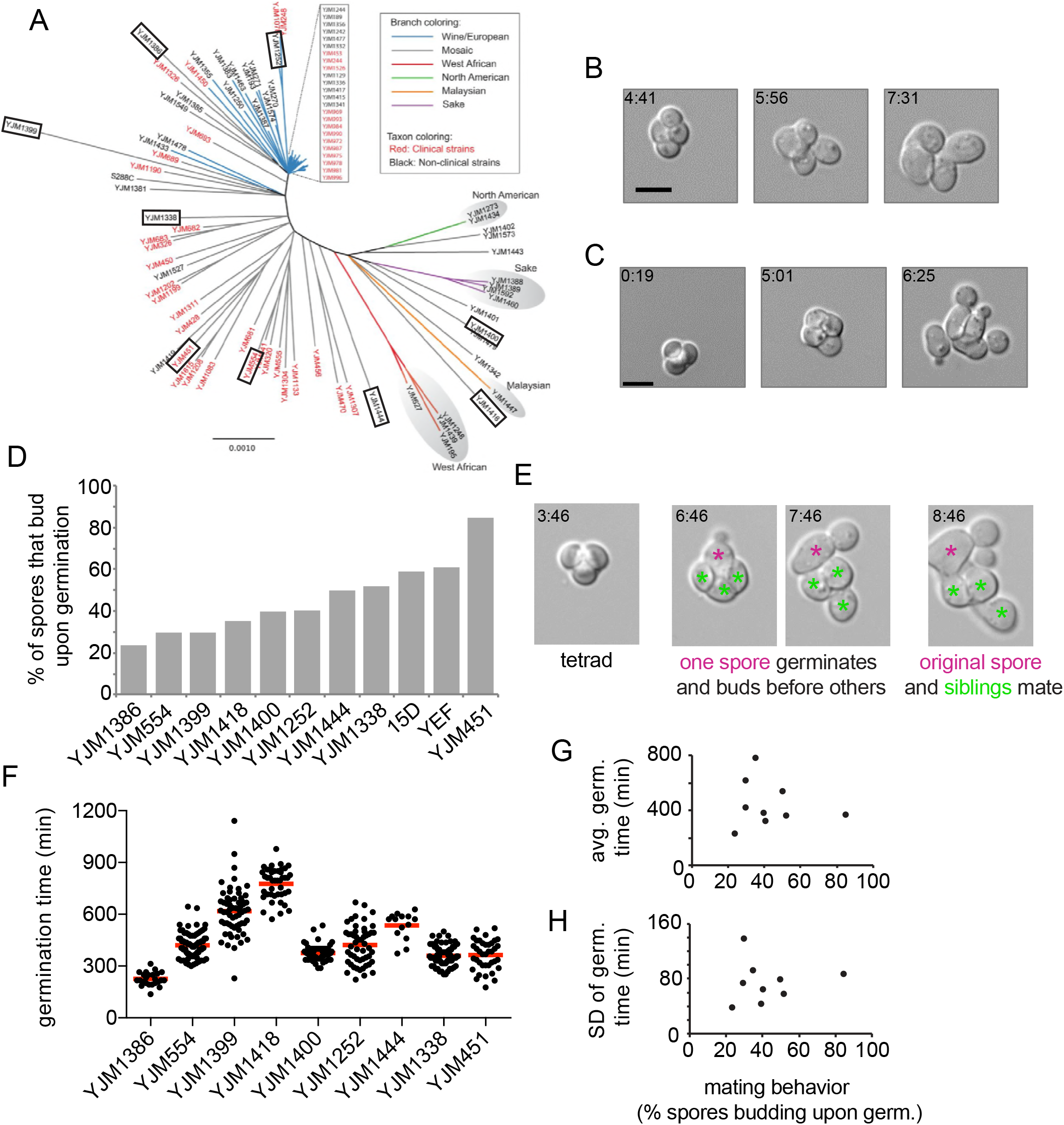
Intratetrad mating behavior following germination. (A) Genetically diverse wild strains used in this study. Image adapted from Strope et al, 2015. (B) Germinating tetrad spores from YJM1418 demonstrating intratetrad mating where all four spores mate (Time after plating on rich media indicated in h:min). (C) Germinating tetrad spores from YJM1338 demonstrating all four spores entering cell cycle and producing buds rather than mating. (D) Quantification of mating behavior upon germination (N>36 spores per strain). (E) Germinating tetrad spores from YJM1418 demonstrating one spore germinating and producing a bud prior to the germination of other spores in same tetrad. Fourth panel shows the original early-germinator mating with an intratetrad partner in its second cell cycle as well as the other two spores mating. (F) Tetrad spores were separated from one another and then placed on an agarose slab containing glucose. Germination time was defined as the time between exposure to glucose and the first time point with a visible bud (N>14 for each strain). (G) and (H) Relation between mating behavior from (B) and average germination time (G) or inter-spore variability in the germination time (H: assessed using the standard devation, SD).

One possible explanation for the failure of sibling spores to mate following germination is a difference in germination times of the spores. If one spore germinates significantly before its potential mating partners, the first spore would have no available partners until after it had undergone one or more cell cycles, and perhaps haplo-selfing after mating-type switching (Fig. 2E). To assess germination timing in the cell populations, we digested tetrads to yield single spores and imaged them. Germination times were indeed variable, both between spores from the same strain and between strains (Fig. 2F). However, there was no obvious correlation between intratetrad mating and germination time (either average time or variability between spores: Fig. 2G,H). Thus, while asynchrony in germination may lower the efficiency of intratetrad mating, it does not appear to be a dominant factor in explaining the different rates of intratetrad mating between strains.

### Mating behavior of isolated germinating spores

As individual spores might be more prevalent than intact tetrads in fly frass, we next separated spores from each other and imaged the events following germination. With no partner nearby, a germinating spore would be expected haplo-self (Fig. 1A). Indeed, some spores followed this behavior (Fig. 3A), but we also observed formation of microcolonies that deferred mating (Fig. 3B). These findings are consistent with the deferred mating observed with tetrads.

**Figure 3.**
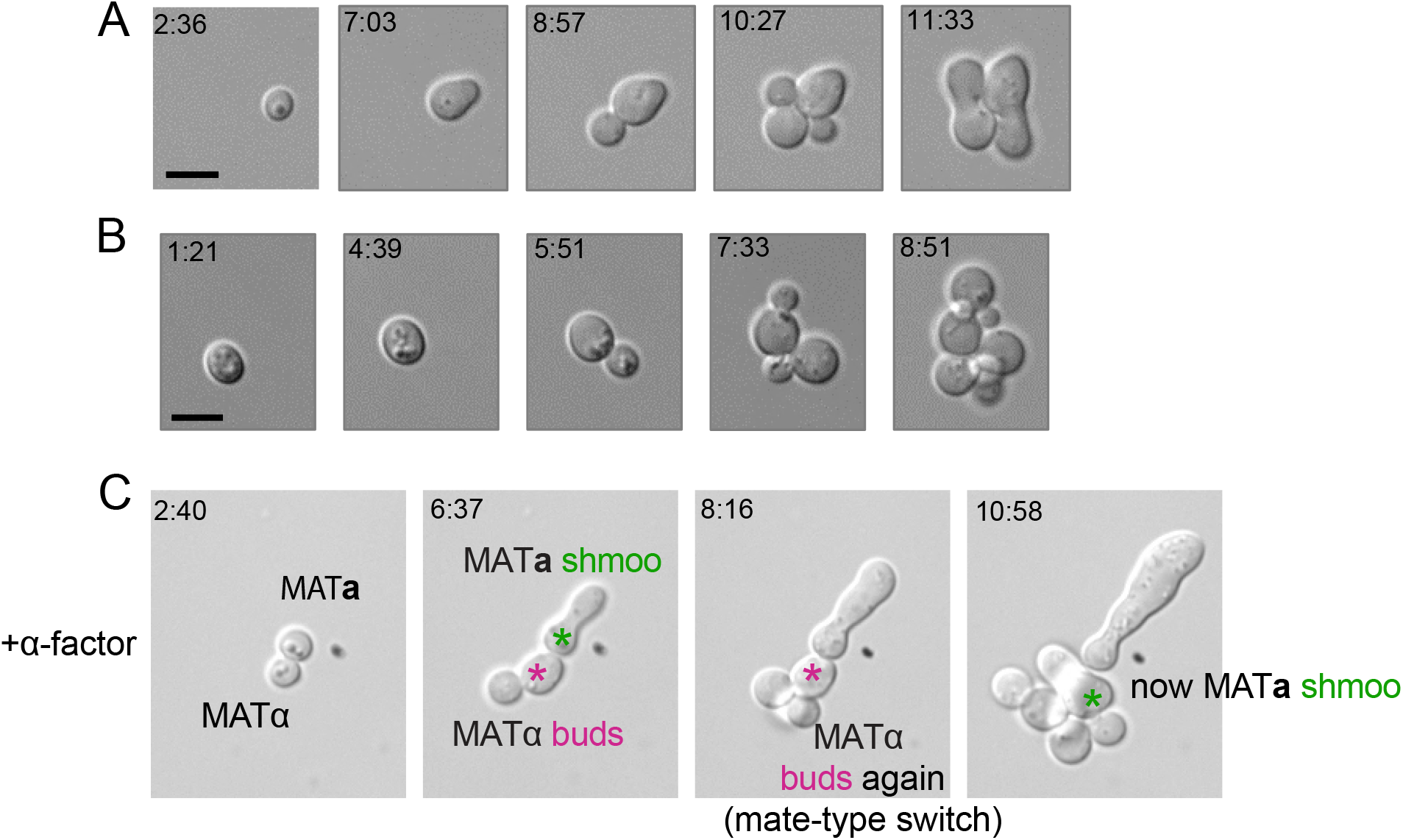
Isolated spore mating behavior following germination. A) Germinating spore from YJM1399 demonstrating haplo-selfing. (B) Germinating spore from YJM451 demonstrating formation of a microcolony through repeated budding. (C) Two isolated spores from YJM1399 germinating in the presence of 2 μM α-factor. The upper spore arrests and forms a shmoo, and the lower spore enters the cell cycle twice and then arrests and forms a shmoo, indicating that it has undergone a mating-type switch.

In principle, deferred mating might result from inefficient mating-type switching, so that opposite mating-type cells do not co-exist until larger microcolonies have formed. To detect mating-type switching from MATα to **a**, we germinated isolated spores in the presence of saturating α-factor. Some spores, presumably MAT**a**, never entered the cell cycle and instead formed a long mating projection. Other germinating spores, presumably MATα, entered the cell cycle and budded. After two budding cycles, these cells arrested and formed a projection, indicating that they had switched mating type (Fig. 3C). In sum, our findings suggest that different strains have inherently different degrees of interest in mating upon germination. We conclude that interest in sex following germination is a variable trait among wild yeast and that many wild mating events are likely to involve cycling haploid cells rather than germinating spores.

### Pheromone sensitivity and production in germinating spores

Potential reasons for a difference in the propensity to mate among wild yeast strains include differences in either the production of pheromones or the reception of pheromones derived from a potential mating partner. For the following set of experiments, we initially focused on strains YJM451 (low propensity to mate; clinical isolate) and YJM1399 (high propensity to mate; cherry tree isolate).

To assess sensitivity to pheromone, spores from each strain were germinated on slabs containing different concentrations of exogenous a-factor. At 2 µM α-factor, 50% of spores (presumably the MAT**a** spores) from both strains germinated, arrested, and formed shmoos (Fig. 4A). At the intermediate concentration of 100 nM α-factor, a mixed response was observed, with some germinating spores elongating (presumably due to a transient G1 arrest) before budding (Fig. 4B). These results suggest that the two strains do not have a drastic difference in pheromone sensitivity.

**Figure 4.**
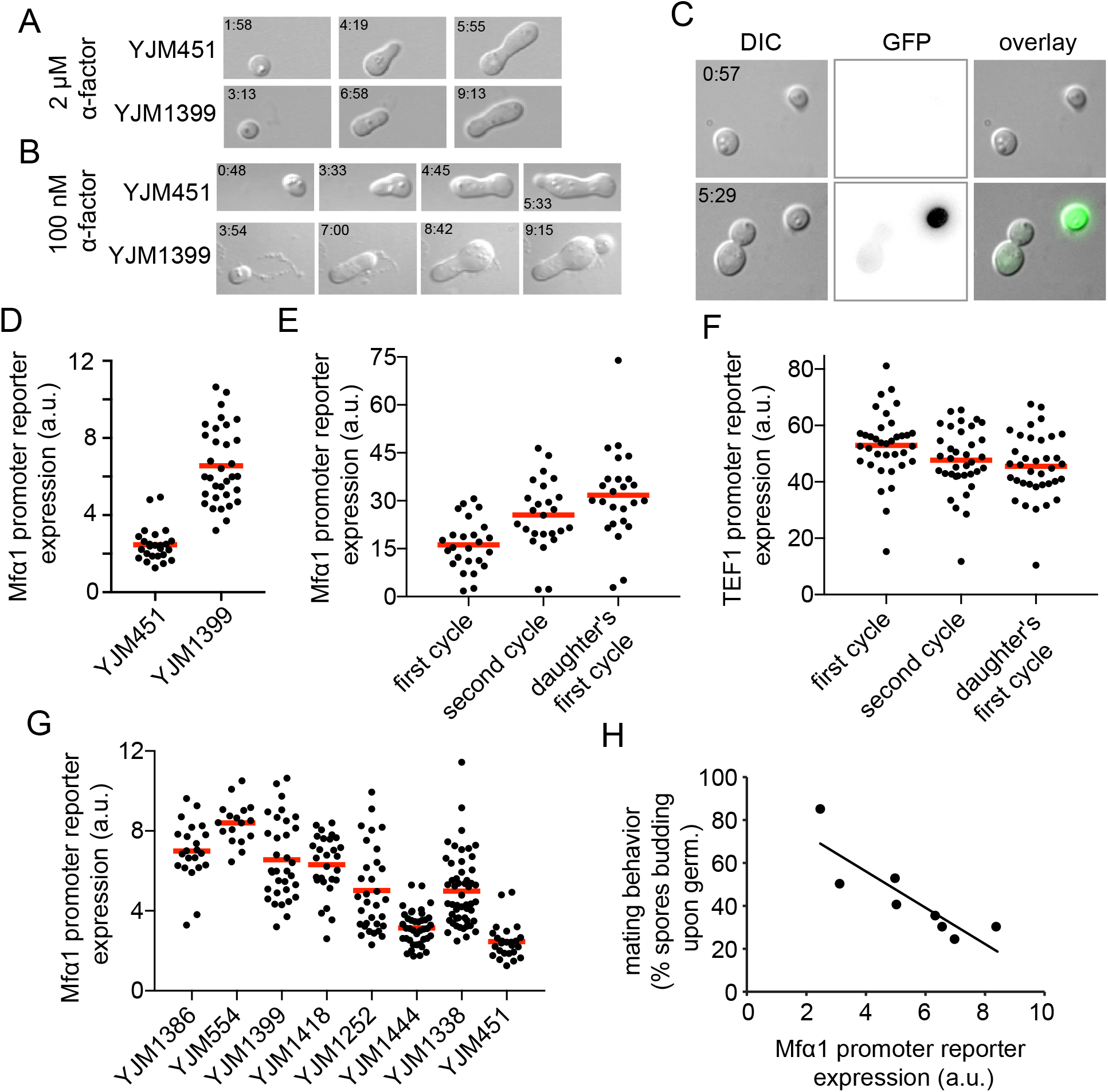
Pheromone sensitivity and pheromone production in germinating spores. Spores of strains YJM451 and YJM1399 were isolated from tetrads and germinated in the presence of 2 μM α-factor (A) or 100 nM α-factor (B). (C) Two isolated spores from DLY20288. The upper-right spore (presumed MATα) shows expression of the Mfα1 reporter following germination whereas the lower-left spore (presumed MAT**a**) does not. (D) Spores from DLY20922 and DLY21020 (containing the Mfα1 reporter integrated into YJM451 and YJM1399) were isolated from tetrads and imaged during germination. Expression of the Mfα1 reporter was determined just prior to bud emergence (see materials and methods). (E) Spores from DLY22729 carrying the Mfα1 reporter were treated as in (D) and expression was determined just prior to bud emergence during the first cell cycle, the second cell cycle, and the first cell cycle of the daughter. The second cell cycle and the daughter signals were each significantly higher than the first cell cycle (t-test, p<0.002). (F) Spores from DLY18930 carrying the TEF1 reporter were treated as in (E). (G) Spores from the indicated wild strains carrying the Mfα1 reporter were treated as in (D) (note that data in D is also included in G). (H) Correlation between the % of spores that bud following germination (from Fig. 2D) and the averaged Mfα1 reporter expression (from Fig. 4F) (Pearson’s correlation, R^2^ = 0.74, p<0.01).

To assess production of pheromone, we integrated a fluorescent reporter at the MFα1 locus (replacing the ORF for pheromone production). As expected, about 50% of germinating spores (presumably MATa) expressed the reporter (Fig. 4C), and we quantified the reporter intensity following germination, 8 min prior to detection of the first bud. Reporter expression was significantly higher for strain YJM1399 than for strain YJM451 (Fig. 4D). Thus, the higher propensity of strain YJM1399 to mate may be due, at least in part, to the higher transcription rate of MFα1.

Interestingly, we found that in our lab strain, expression of the MFα1 reporter was higher in the second than in the first cell cycle following germination (Fig. 4E). Moreover, daughter cells made by the germinated mother expressed higher levels of the reporter in their first cell cycle than the germinating mothers had (Fig. 4E). In all cases, the reporter expression level was measured at the timepoint prior to bud emergence. In contrast to MFα1, expression from a control *TEF1* promoter was similar in the first and second cell cycles following germination, and in daughter cells (Fig. 4F). These findings suggest that pheromone production ramps up over the first two (and perhaps more) cycles following germination. If the magnitude and timing of such a ramp-up were variable between strains, then that might lead to the observed variability in the propensity to mate immediately following germination.

To assess the degree to which mating propensity is correlated with MFα1 expression, we integrated the reporter at the MFα1 locus for the entire set of strains. MFα1 expression showed significant variability between strains (Fig. 4G) and was well correlated with intratetrad mating propensity (R^2^=0.74, p<0.01, Fig. 4H). This correlation suggests that variability in the amount of pheromone produced at the time of germination may explain some of the variability in the mating behavior of wild strains.

### Basis for variability in pheromone production in wild strains

A potential explanation for variable pheromone production could be variability in the synthetic capacity of germinating spores. In particular, there was considerable variation in the size of the spores, and we speculated that larger spores might produce more pheromone. However, we did not detect any correlation between cell size and MFα1 reporter expression, either within or between strains (Fig. 5A). Thus, it seems likely that variable MFα1 expression is due to cis- or trans-acting factors affecting the promoter. To test whether trans-acting factors were the dominant contributor to the variable expression, we integrated a reporter driven by the MFα1 promoter from a lab strain into each wild strain. The different wild strains exhibited differences in reporter expression at the time of germination (Fig. 5B), indicating that trans-acting factors vary between the strains. Surprisingly, comparison of MFα1 expression from the transgene and from the endogenous locus revealed little correlation (Fig. 5C), suggesting that cis-acting alterations also play a role in MFα1 expression. Examination of the genome sequences revealed several polymorphisms in the MFα1 promoter (Fig. S1), consistent with that possibility.

**Figure 5.**
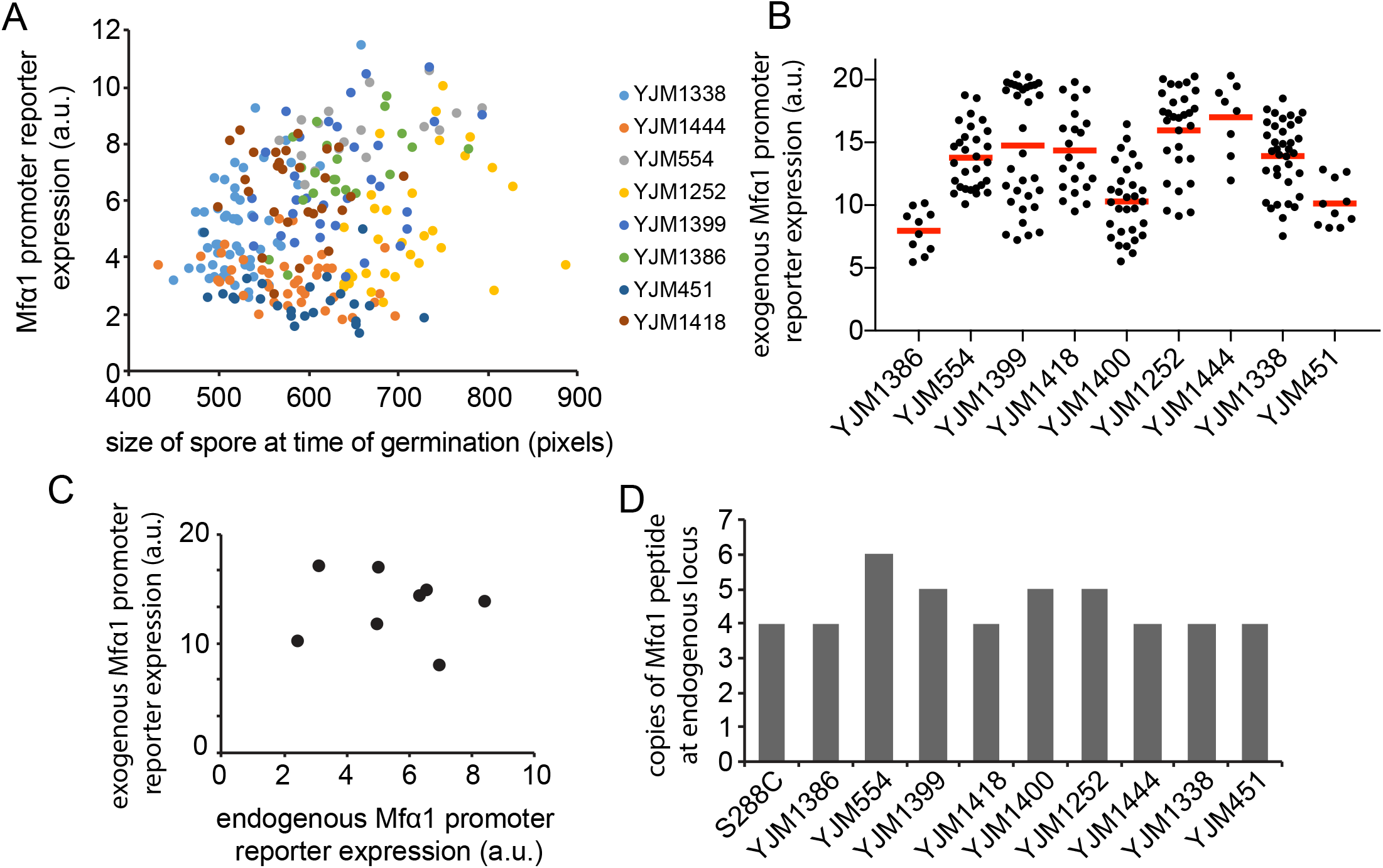
Basis for differences in pheromone production between strains. (A) Relation between spore size and Mfα1 reporter expression in germinating spores. Each dot is one cell, color-coded by strain. (B) Mfα1 reporter driven by an exogenous lab strain Mfα1 promoter shows variable expression in different strains. (C) Relation between average endogenous Mfα1 reporter expression and average exogenous Mfα1 reporter expression. (D) Copy number of Mfα1 peptides at the endogenous locus of the different wild strains (Strope et al., 2015).

We note that pheromone production depends on more than just MFα1 transcription. There are two genes encoding α-factor (MFα1 and MFa2), and pheromone production involves proteolytic processing of the primary protein product, which encodes 4 copies of the pheromone peptide in lab strains. We found that the number of pheromone peptide copies in MFα1 varied (up to 6 copies) among the wild strains we examined (Fig. 5D) (Strope et al., 2015), presumably contributing to variation in pheromone secretion. In sum, these data suggest that pheromone expression is variable in wild strains, due to both cis- and trans-acting differences between strains.

## Discussion

The yeast mating pathway is one of the best-understood information-processing systems in eukaryotic biology. Recent work has highlighted the decision-making capabilities of the pheromone response pathway, revealing behaviors that have been interpreted as optimizing the mating potential of the cells under imagined scenarios motivated by consideration of lab strains that are heterothallic (incapable of mating-type switching and therefore haplo-selfing)(Banderas et al., 2016; Caudron and Barral, 2013; Doncic et al., 2015; Erdman and Snyder, 2001; Hao et al., 2008; Paliwal et al., 2007). However, yeast in the wild are homothallic, so mating scenarios in the wild may differ significantly. Thus, it is important to understand the likely situations that the information processing capabilities of this pathway were evolved to accommodate. Our primary finding is that many yeast strains derived from wild isolates show a low propensity to mate upon germination. This behavior has significant implications.

One consequence of the delayed interest in sex is that germinating spores would undergo a few cell cycles as haploids before mating. This provides an opportunity for natural selection to purge deleterious mutations and enrich for beneficial mutations.

Another consequence of delayed interest in sex is to reduce the reproductive isolation that might arise as a result of unsynchronous germination. Our work suggests that in many wild strains, there is considerable spore-to-spore variability in germination time (e.g. strain YJM1399 in Fig. 2F). Differences in germination time have been proposed to promote reproductive isolation, leading to speciation (Murphy and Zeyl, 2012). However, delayed propensity to mate upon germination means that mating would often occur between haploid cells in microcolonies, rather than between germinating spores themselves. That would allow opportunities to mate between progeny of spores that germinated at different times.

Three main scenarios have been envisaged for mating of wild yeast: haplo-selfing, intratetrad mating, and outbreeding (Knop, 2006; Liti, 2015). If a spore were to germinate in the absence of a nearby partner, then mother-specific mating-type switching would lead to haplo-selfing, generating homozygous diploids. If spores in an intact tetrad were to germinate synchronously, then they would mate without intervening mitoses to generate partially homozygous diploids (intratetrad mating). And if a spore were to germinate alongside a spore from an unrelated tetrad, the germinating spores could mate to form a fully heterozygous diploid (outbreeding). The conditions under which outbreeding would occur are not well understood, but high-density plating of tetrads on germination-promoting media revealed that germinating spores often mated with partners from neighboring tetrads (Murphy and Zeyl, 2010). Moreover, passage through the gut of Drosophila (an avid eater of yeast) can break up tetrads during digestion, depositing unrelated spores in the frass (Reuter et al., 2007).

The high levels of heterozygosity observed in some wild diploid yeasts (Kelly et al., 2012; Liti et al., 2009; Magwene et al., 2011; Muller and McCusker, 2009; Strope et al., 2015) suggest that outbreeding occurs with appreciable frequency, particularly in human-associated environments. Delayed interest in sex could lead to a greater frequency of outbreeding. If germinating spores were the mating entity, then unrelated spores would have to find themselves in very close proximity (and germinate synchronously) in order to mate. However, if a haploid microcolony is formed by one or both spores, cells at the periphery of each microcolony would have the opportunity to mate to potentially genetically different partners, yielding outbred diploids.

## Acknowledgements

We thank P. Magwene for the wild yeast strains and for helpful advice throughout the project. We thank N. Buchler, B. Errede, A. Gladfelter, D. Kiehart, P. Magwene, J. McCusker, and members of the D.J.L. lab for thoughtful discussions and critical readings of the manuscript. Special thanks to the D. Fox lab for supplying flies and helping with fly techniques and to D. McClure for help with data analysis. This work was funded by NIH/NIGMS grants GM103870 and GM122488 to D.J.L.

## Author Contributions

A.W.M., K.J., and D.J.L. designed and analyzed experiments; A.W.M. and K.J. performed experiments; T.R.Z. generated plasmids and yeast strains; and A.W.M. and D.J.L. wrote the paper with contributions from K.J.

## Materials and Methods

### Yeast strains and plasmids

Standard molecular genetic procedures were used for strain construction. Yeast strains, plasmids, and construction details are listed in the Supplement. Wild yeast strains were originally isolated and characterized in Strope et al, 2015.

### Sporulation conditions and spore isolation

Prior to sporulation, yeast were grown to near saturation in liquid YEPD rich media at 30°C. Approximately 8 ml of saturated culture were washed briefly with water and resuspended in 8 ml of 2% KAC media (2% potassium acetate supplemented with 0.004% adenine, 0.003% histidine, 0.004% uracil, 0.008% leucine, 0.003% tryptophan). Cultures were kept at 30°C for 10 days, then moved to 4°C.

To separate spores from tetrads, 1 ml of sporulated cells were washed with water, resuspended in 20-30 μl of lyticase solution (2.72% lyticase powder, 1 M sorbitol, 100 mM PIPES pH 6.5), and incubated for 20 min at room temperature. Spores were then resuspended in 1 ml of water and sonicated at 50% amplitude for approximately 20 s. Samples were heated at 55°C for 30 min to kill any remaining diploid cells, then stored at 4°C.

### Feeding yeast to flies

1 ml of sporulated yeast cultures were washed with water and resuspended in the residual water. 5 μl of this paste was pipetted on a 30 mm grape plate, and 50 w1118 flies were then incubated in the plate for 6 h. 15 flies were then transferred to a 10 cm YEPD plate or a YEPD plate supplemented with clonNAT and kanamycin for 1 h. After removing the flies, excrement spots were noted, and the next day, the spots from the YEPD plate were streaked onto a YEPD plate with both drugs.

### Filter mating assay

Quantitative, filter-based mating assays were essentially as performed in Hartwell, 1973. 5*10^5^ spores (tetrads were counted as 4) from each genotype were mixed and filtered onto triplicate nitrocellulose filters, which were placed on a YEPD agarose plate. After 4 h, cells were washed in PBS and sonicated in 50 ml conical tubes. 150 cells from each filter were plated in triplicate on YEPD plates supplemented with clonNAT or kanamycin, and 2000 cells were plated on YEPD plates with both drugs. Outcross frequency was calculated as the percentage of recovered cells that were resistant to both drugs (NAT^R^KAN^R^ / (NAT^R^ + KAN^R^ – NAT^R^KAN^R^)).

### Live cell imaging

100-200 μl of spores (see sporulation conditions and isolation section) were washed once with water, then resuspended in approximately 8 μl of media (2% dextrose in complete synthetic media). 0.75 μl of cells were mounted onto a 2% agarose slab with the same media, sealed with petroleum jelly, and kept in a humidity chamber at 30°C prior to imaging.

Live cell imaging was performed on a Zeiss Axio Observer essentially as described in Howell et al, 2012. DIC images were acquired every 3 min using the autofocus function in the MetaMorph software. If fluorescence images were also being acquired, then DIC images with autofocus were acquired every 8 min, and fluorescence images were taken every 16 min. GFP images were acquired with 800 EM gain, 200 ms exposure and 20 z-steps that were 0.42 μm apart. GFP quantification of MFα1 reporters was performed on the images acquired at the time point just prior to visualizing budding. The fluorescence zstack was maximum projected and the mean fluorescence value was recorded. For experiments using each strain’s endogenous MFα1 promoter or TEF1 promoter, mean fluorescence values were normalized against Bem1-GFP fluorescence in DLY19805 that was germinated and imaged concurrently.

## References

Alvaro, C.G., and Thorner, J. (2016). Heterotrimeric G protein-coupled receptor signaling in yeast mating pheromone response. J. Biol. Chem. 291, 7785–7798.

Arkowitz, R.A. (2009). Chemical gradients and chemotropism in yeast. Cold Spring Harb. Perspect. Biol. 1, a001958.

Atay, O., and Skotheim, J.M. (2017). Spatial and temporal signal processing and decision making by MAPK pathways. J. Cell Biol. 216, 317–330.

Banderas, A., Koltai, M., Anders, A., and Sourjik, V. (2016). Sensory input attenuation allows predictive sexual response in yeast. Nat. Commun. 7, 1–9.

Botstein, D., and Fink, G.R. (2011). Yeast: An experimental organism for 21st century biology. Genetics 189, 695–704.

Botstein, D., Chervitz, S.A., and Cherry, J.M. (1997). Yeast as a model organism. Science. 277, 1259–1260.

Boynton, P.J., and Greig, D. (2014). The ecology and evolution of non-domesticated Saccharomyces species. Yeast 31, 449–462.

Caudron, F., and Barral, Y. (2013). A super-assembly of Whi3 encodes memory of deceptive encounters by single cells during yeast courtship. Cell 155, 1244–1257.

Coluccio, A.E., Rodriguez, R.K., Kernan, M.J., and Neiman, A.M. (2008). The yeast spore wall enables spores to survive passage through the digestive tract of Drosophila. PLoS One 3, 1–7.

Doncic, A., Atay, O., Valk, E., Grande, A., Bush, A., Vasen, G., Colman-Lerner, A., Loog, M., and Skotheim, J.M. (2015). Compartmentalization of a Bistable Switch Enables Memory to Cross a Feedback-Driven Transition. Cell 160, 1182–1195.

Erdman, S., and Snyder, M. (2001). A filamentous growth response mediated by the yeast mating pathway. Genetics 159, 919–928.

Gilbert, D.G. (1980). Dispersal of yeasts and bacteria by Drosophilia in a temperate forest. Oecologia 46, 135–137.

Haber, J.E. (2012). Mating-type genes and MAT switching in Saccharomyces cerevisiae. Genetics 191, 33–64.

Hao, N., Nayak, S., Behar, M., Shanks, R.H., Nagiec, M.J., Errede, B., Hasty, J., Elston, T.C., and Dohlman, H.G. (2008). Regulation of cell signaling dynamics by the protein kinase-scaffold Ste5. Mol Cell 30, 649–656.

Hartwell, L.H. (1973). Synchronization of haploid yeast cell cycles, a prelude to conjugation. Exp Cell Res 76, 111–117.

Herman, P., and Rine, J. (1997). Yeast spore germination: a requirement for Ras protein activity during re-entry into the cell cycle. EMBO J. 16, 6171–6181.

Joseph-Strauss, D., Zenvirth, D., Simchen, G., and Barkai, N. (2007). Spore germination in Saccharomyces cerevisiae: global gene expression patterns and cell cycle landmarks. Genome Biol. 8, R241.

Kelly, A.C., Shewmaker, F.P., Kryndushkin, D., and Wickner, R.B. (2012). Sex, prions, and plasmids in yeast. Proc. Natl. Acad. Sci. 109, E2683–E2690.

Knop, M. (2006). Evolution of the hemiascomycete yeasts: on life styles and the importance of inbreeding. Bioessays 28, 696–708.

Liti, G. (2015). The fascinating and secret wild life of the budding yeast S. cerevisiae. Elife 4, 1–9.

Liti, G., Carter, D.M., Moses, A.M., Warringer, J., Parts, L., James, S.A., Davey, R.P., Roberts, I.N., Burt, A., Koufopanou, V., et al. (2009). Population genomics of domestic and wild yeasts. Nature 458, 337–341.

Magwene, P.M., Kayıkçı, Ö., Granek, J. a, Reininga, J.M., Scholl, Z., and Murray, D. (2011). Outcrossing, mitotic recombination, and life-history trade-offs shape genome evolution in Saccharomyces cerevisiae. Proc. Natl. Acad. Sci. U. S. A. 108, 1987–1992.

Merlini, L., Dudin, O., and Martin, S.G. (2013). Mate and fuse: how yeast cells do it. Open Biol. 3, 1–13.

Michaelis, S., and Barrowman, J. (2012). Biogenesis of the Saccharomyces cerevisiae Pheromone a-Factor, from Yeast Mating to Human Disease. Microbiol. Mol. Biol. Rev. 76, 626–651.

Muller, L. a H., and McCusker, J.H. (2009). Microsatellite analysis of genetic diversity among clinical and nonclinical Saccharomyces cerevisiae isolates suggests heterozygote advantage in clinical environments. Mol. Ecol. 18, 2779–2786.

Murphy, H. a., and Zeyl, C.W. (2010). Yeast sex: surprisingly high rates of outcrossing between asci. PLoS One 5, 19–21.

Murphy, H. a., and Zeyl, C.W. (2012). Prezygotic isolation between saccharomyces cerevisiae and saccharomyces paradoxus through differences in mating speed and germination timing. Evolution (N. Y). 66, 1196–1209.

Neiman, A.M. (2011). Sporulation in the budding yeast Saccharomyces cerevisiae. Genetics 189, 737–765.

Paliwal, S., Iglesias, P.A., Campbell, K., Hilioti, Z., Groisman, A., and Levchenko, A. (2007). MAPK-mediated bimodal gene expression and adaptive gradient sensing in yeast. Nature 446, 46–51.

Reuter, M., Bell, G., and Greig, D. (2007). Increased outbreeding in yeast in response to dispersal by an insect vector. Curr. Biol. 17, 81–83.

Stefanini, I., Dapporto, L., Berná, L., Polsinelli, M., Turillazzi, S., and Cavalieri, D. (2016). Social wasps are a Saccharomyces mating nest. Proc. Natl. Acad. Sci. 113, 2247–2251.

Strope, P.K., Skelly, D. a, Kozmin, S.G., Mahadevan, G., Stone, E. a, Magwene, P.M., Dietrich, F.S., and McCusker, J.H. (2015). The 100-genomes strains, an S. cerevisiae resource that illuminates its natural phenotypic and genotypic variation and emergence as an opportunistic pathogen. Genome Res. 25, 762–774.

Taxis, C., Keller, P., Kavagiou, Z., Jensen, L.J., Colombelli, J., Bork, P., Stelzer, E.H.K., and Knop, M. (2005). Spore number control and breeding in Saccharomyces cerevisiae: a key role for a self-organizing system. J. Cell Biol. 171, 627–640.

